# Integrating Ambient Mass Spectrometry Imaging and Spatial Transcriptomics on the Same Cancer Tissues to Identify Gene-Metabolite Correlations

**DOI:** 10.1101/2024.12.20.626670

**Authors:** Trevor M. Godfrey, Yasmin Shanneik, Wanqiu Zhang, Thao Tran, Nico Verbeeck, Nathan H. Patterson, Faith E. Jackobs, Chandandeep Nagi, Maheshwari Ramineni, Livia S. Eberlin

## Abstract

The combination of spatial metabolomics and spatial transcriptomics can reveal powerful connections between gene expression and metabolism in heterogeneous tissues, but section-to-section variability can convolute data integration. We present a novel method combining Desorption Electrospray Ionization Mass Spectrometry Imaging spatial metabolomics and spatial transcriptomics on the same tissue sections. We demonstrate this workflow on human breast and lung cancer tissues identifying correlations between metabolites and cancer-related RNA transcripts.

## Main

The study of complex biological tissues necessitates the use of spatial omics techniques to better understand heterogeneous microenvironments. Visium Spatial Transcriptomics (VST) has emerged as a predominant technique to map the spatial heterogeneity of gene expression. Mass spectrometry imaging (MSI) is the leading method to characterize the distribution of metabolites and lipids. Powerful contributions to cancer biology have been made possible using VST and MSI to map molecular changes within the tumor microenvironment.^1,2^ However, when used separately, MSI and VST offer a limited view into specific classes of molecules. Understanding the complex interplay between gene expression and metabolic phenotypes requires precise integration of spatial transcriptomics (ST) and spatial metabolomics (SM) datasets. This can be accomplished by performing MSI and VST workflows on adjacent serial sections.^3–6^ Yet, biological and technical variability between serial tissue sections leads to challenges and ambiguity in data alignment and interpretation. While computational methods have been developed to address co-registration challenges, obtaining data from the same tissue section is still the ideal approach.

In this study, we describe the development of a method that integrates desorption electrospray ionization (DESI)-MSI and VST workflows on the same human cancer tissue sections. We have called this multi-modal method Desium (a portmanteau of DESI and Visium). DESI is a MSI technique performed in ambient conditions to analyze the spatial distribution of molecules directly from a tissue section without tissue damage.^7^ A protocol combining matrix-assisted laser desorption ionization (MALDI)-MSI and VST on the same tissue section has been described by Vicari *et al*.^8^ The authors demonstrated the compatibility of four MALDI matrices with VST in separate experiments targeting either lipids or metabolites. A semi-targeted analysis of neurotransmitters was reported in conjunction with VST results on mouse and human Parkinson’s Disease tissues. Unlike MALDI-MSI, DESI-MSI requires no sample preparation or matrix application. With DESI-MSI, lipids and metabolites are detected in a single experimental run, allowing for the collection of rich untargeted data. Additionally, the same tissue section may be reimaged in different ionization modes or in other multi-modal analyses.^7,9,10^ Ease-of-use, speed, and non-destructivity make DESI-MSI an attractive technique to perform large, untargeted, multi-modal MSI studies. Yet, DESI-MSI has not been shown to be compatible with RNA analyses, such as VST. In this study, we show that performing DESI-MSI before VST does not impact RNA quality, allowing for the easy and rapid acquisition of MSI data prior to collecting VST data on the same tissue section.

The Desium workflow we developed consists of tissue sectioning, DESI-MSI at 100 μm spatial resolution, hematoxylin and eosin (H&E) staining, and 100 μm VST analyses performed sequentially on the same tissue section (**Fig. 1a**). To assess whether DESI-MSI affects RNA quality, we measured the RNA Integrity Number (RIN) on tissue sections from five mice brains after DESI-MSI. A RIN of 4 is the minimum value required to perform the VST CytAssist protocol. The RIN of each replicate ranged from 8.6-9.1, indicating exceptional RNA quality after DESI-MS imaging analysis (**Extended Data** Figs. 1a-1b).

**Figure 1:**
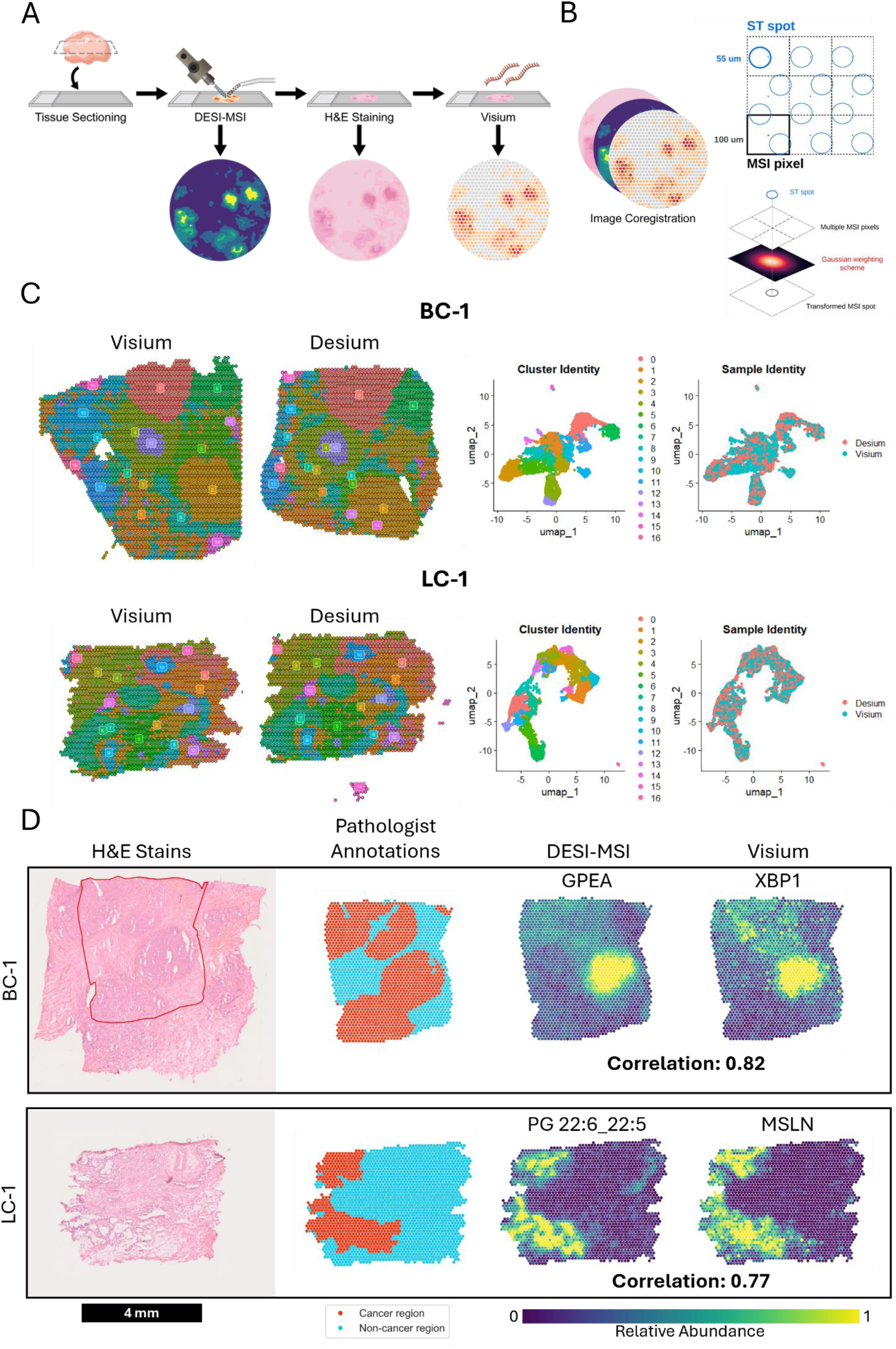
Desium, a spatial multiomic workflow to investigate spatial metabolomics and transcriptomics on the same tissue section. **a,** The Desium workflow–fresh-frozen tissues are sectioned and mounted onto glass slides and DESI-MSI is performed with 100% methanol spray-solvent. Tissues are H&E stained and imaged with bright-field microscopy. The VST CytAssist protocol for fresh frozen tissue is then employed. **b,** DESI-MSI and VST data are aligned to post-MSI H&E and Visium CytAssist H&E images with a manual nonrigid coregistration based on histological and molecular landmarks and automatic linear co-registration respectively; H&E images and corresponding data are aligned with manual non-rigid coregistration. MSI data was converted into VST spot coordinates by applying a Gaussian granularity matching algorithm. **c,** UMAP clustering of VST data from Visium only and Desium workflows display quasi-identical clustering for both breast (breast cancer sample BC-1) and lung (lung cancer sample LC-1) cancer tissues in real and in UMAP space. **d,** The highest top individual lipid/metabolite to gene Pearson correlations for BC-1 and LC-1 aligned with histological features that pathologists identified as cancer regions. Due to tissue folding that occurred on BC-1 during the CytAssist step of the VST protocol, a region of interest outlined in red was selected so that only tissue unaffected by folds was considered in subsequent coregistration and analysis. GPEA: glycerylphosphorylethanolamine, PG: Phosphatidylglycerol. Lipid fatty acid chain notation: (fatty acid chain one)_(fatty acid chain two), (# of carbons in FA chain):(# of double bonds in chain).

Next, we analyzed six human cancer tissues (3 invasive lobular breast cancer and 3 adenocarcinoma lung cancer) using the Desium workflow. For two samples (1 breast, 1 lung), a serial section was analyzed using only the standard VST CytAssist protocol as a control. A variety of peaks previously identified as lipids and metabolites were observed in the MS data for all tissue sections analyzed, typical of what has been previously reported for untargeted DESI-MSI analyses of human breast and lung cancer tissues (**Extended Data** Fig. 2a).^11,12^ Tentative identifications for 159 of the molecules detected from all tissues including 63 that were confirmed with MS2 experiments are listed in supplementary table 2. Importantly, when comparing the sequencing results between Desium and standard VST protocol using the unique molecular identifier (UMI) counts/gene, we found a correlation coefficient of 1 for both lung and breast data, indicating that there are little to no alterations in the measurement of gene expression due to performing DESI-MSI prior to VST (**Extended Data** Fig. 1c). We also observed similar distributions of UMIs/spot and genes/spot between workflows (**Extended Data** Figs. 1d-1e). Uniform Manifold Approximation and Projection (UMAP) analysis revealed the same spatial clusters in tissues analyzed with the Desium workflow and VST alone (**Fig. 1c**). These results confirm that performing DESI-MSI on a tissue prior to VST has little to no impact on VST data quality.

Because DESI-MSI and VST were performed on the same tissue section, we were able to co-register H&E images, DESI-MSI, and VST data to a common coordinate system with a Gaussian weighting scheme to transform MSI pixels and pathologist annotations into a Visium staggered spot format (**Fig. 1b**). We calculated Pearson correlation scores between every gene and metabolite for each tissue and tissue class to identify potential RNA transcripts and lipids/metabolites relationships. We found 6,075 individual transcript-metabolite correlations between 980 transcripts and 133 metabolites with a correlation score above 0.5 or below -0.5. From these, 103 transcript-metabolite correlations above 0.7 were identified among all samples. A list of these correlations is found in supplementary table 4. The top sample-specific correlations are shown in **Fig. 1d** and **Extended Data** Fig. 2b. Notably, a 0.82 correlation between glycerophosphorylethanolamine (GPEA) and XBP1 was observed in breast cancer tissue BC-1 (**Fig. 1d)**. XBP1 is a transcription factor known to promote breast cancer and the transcription of genes involved in lipid biosynthesis.^13^ GPEA, a product of lipid catabolism, has been observed at higher levels in other cancers.^14^ In lung cancer sample LC-1, we found a correlation of 0.77 between phosphatidylglycerol PG 22:6_22:5 and MSLN, a lipid anchored differentiation antigen present in 70% of lung cancers.^15^

The top lipid/gene correlations across tissue classes were lower than for any individual tissue, possibly due to inter-tumor heterogeneity, but still revealed interesting trends for each tissue class. In breast cancer tissues, we observed a 0.57 average correlation score between phosphatidyl-ethanolamine (PE) 18:2_18:0 and MUC1 (**Extended Data** Fig. 3c). In lung cancer samples, we observed a 0.42 correlation score between a PE chain with a different fatty acid composition, PE 18:1_20:4, and MUC1. MUC1 is a gene that encodes a transmembrane glycoprotein known to increase lipid metabolism and transportation in multiple epithelial cancers, including breast and lung cancer.^16^ A list of mean correlations between transcripts and identified metabolites is found in supplementary table 6.

To visualize overall trends from the individual gene and lipid/metabolite correlations identified in this workflow we utilized non-negative matrix factorization (NMF) component analysis and calculated Pearson correlation scores between the DESI-MSI, VST, and histological regions of interest (**Fig. 2** and **Extended Data** Figs. 4-10). DESI-MSI and VST NMF components display high spatial correlations for many tissue regions, revealing fascinating trends in tumor heterogeneity that could not be seen by histopathology alone. For example, clear intra-tumor heterogeneity was seen in breast cancer sample BC-1 where NMF analysis revealed three distinct molecular distributions for both DESI-MSI and VST data across three tumor regions that were assessed as identical by histopathology alone (**Fig. 2d**). These distinct tumor regions were observed in both DESI-MSI and VST datasets and were highly correlated between datasets. In lung cancer sample LC-1, NMF analysis reveals molecular patterns corresponding to regions of tumor and lung airways (**Fig. 2h**).

**Figure 2:**
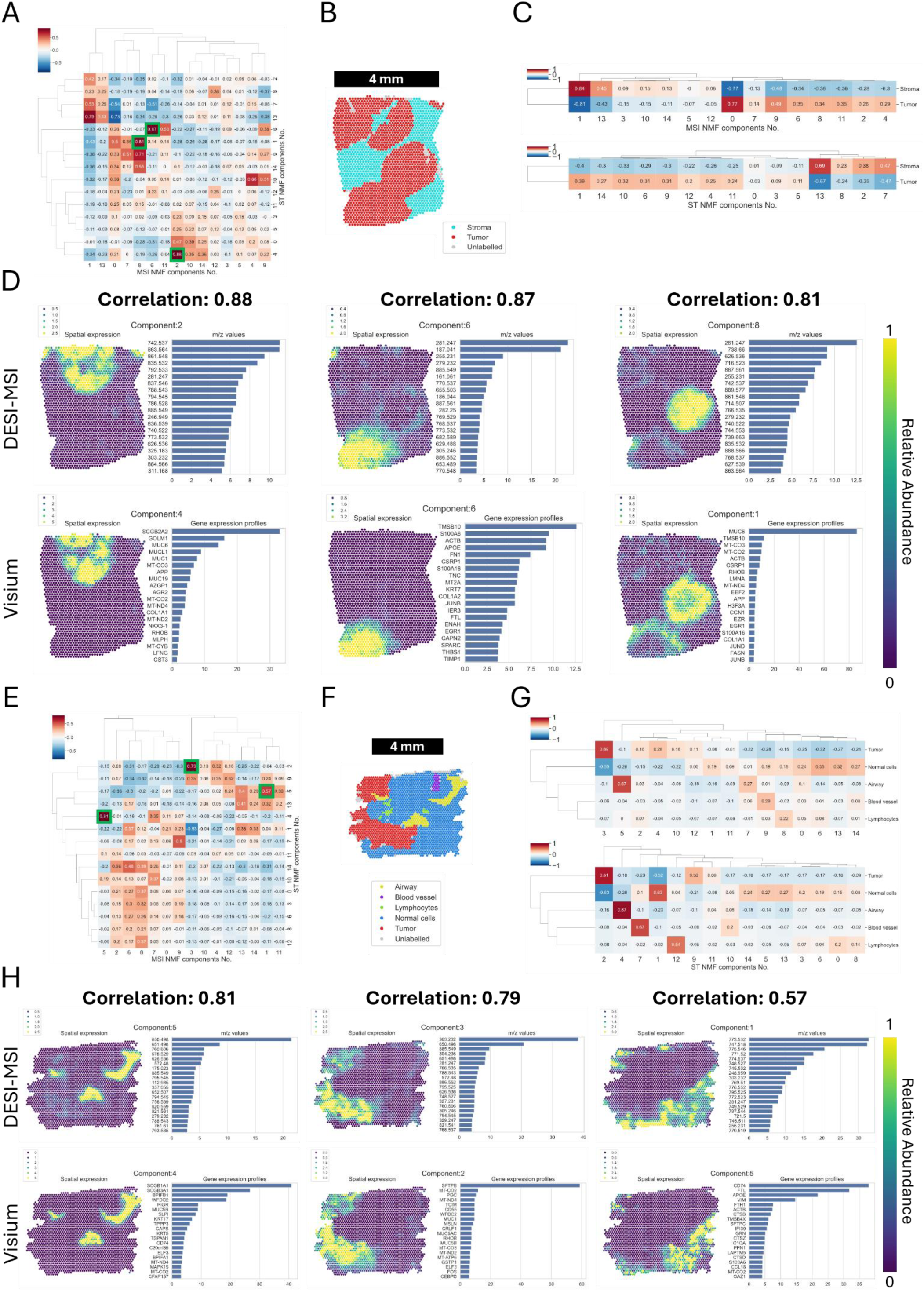
Multidimensional NMF Analysis of Human Breast and Lung Cancer Desium Data. **a**, Heat map showing Pearson correlation scores between DESI-MSI and VST NMF components for breast cancer sample BC-1. The three highest correlations are highlighted with a green box. **b,** Pathologist annotations of BC-1 transformed into Visium spot format. **c,** Heatmaps showing the Pearson correlation score of each NMF component with pathologist identified histological features. **d,** NMF component analysis of MSI and VST data display highlight clear intratumor heterogeneity in BC-1. Strong correlations between these subregions are seen between DESI-MSI and VST data. The top twenty contributors for each NMF component are shown to the right of each image. **e,** Heat map showing Pearson correlation scores between DESI-MSI and VST NMF components for Lung cancer sample LC-1. The three highest correlations are highlighted with a green box. **f,** Pathologist annotations of LC-1 transformed into Visium spot format. **g,** Heatmaps showing the Pearson correlation score of each NMF component with pathologist histological features. **h,** NMF component analysis of MSI and VST data highlight clear molecular patterns between cancer, airways, and normal cells in LC-1. Strong correlations between these subregions are seen between DESI-MSI and VST data. The top twenty contributors for each NMF component are shown to the right of each image.

In summary, we present a workflow that enables simultaneous analysis of metabolites, lipids, and gene expression all on the same tissue. Our results show that RNA is still intact after performing DESI-MSI on tissues, indicating that DESI-MSI may also be compatible with other RNA analysis techniques such as single cell RNA-seq, fluorescence in situ hybridization, and in-situ sequencing. Importantly, integration of DESI-MSI and VST on the same tissue sections at the same spatial resolution allows for more precise alignment of the datasets, eliminates inter-section biological and technical variability, and allows maximum data acquisition from a single tissue.

In this study, unambiguous spatial correlations measured using the Desium workflow identified relationships between lipids, metabolites, and RNA transcripts, and revealed unique patterns of intratumor heterogeneity that were not captured by histologic analysis. Altogether, our study shows that Desium can be a powerful discovery tool to understand the relationship between gene expression and the metabolic phenotype of cells in the complex tumor microenvironment.

## Materials and Methods

### Tissue samples collection and preparation

Five CD-1 (ICR) gender unspecified mouse brain samples were purchased from BioIVT and used to assess RNA degradation after DESI-MSI. Three human invasive lobular breast cancer samples and three human adenocarcinoma lung cancer samples were acquired from the Cooperative Human Tissue Network (CHTN) to test the Desium workflow. Gross pathological evaluations for each tissue can be found in supplemental table 1. Mouse brain samples were assigned labels MB1, MB2, MB3, MB4, and MB5. Human breast cancer samples were assigned labels BC-1, BC-2, and BC-3. Human lung cancer samples were assigned labels LC-1, LC-2, and LC-3. All samples were stored at -80 °C and equilibrated and sectioned in a cryostat (Epredia CryoStar NX50) at temperatures between -10 °C to -15 °C. Mouse brain samples were sectioned at 10 μm thickness. Human breast samples were sectioned at 7 μm thickness. Human lung samples were sectioned at 5 µm thickness. Tissues were thaw mounted on room temperature plus-charged slides (Superfrost Plus 25 x 75 x 1.0 mm, Fisher Scientific) and set on a 37 °C warm plate for 10 seconds to thaw and dry samples. Each thaw mounted tissue was taken directly for DESI-MSI or VST analysis.

### Assessment of RNA degradation post-DESI-MSI

Three serial sections each were collected from the Bregma region of the five mouse brains (MB1-MB5) for RNA degradation assessment. Control serial sections were collected in chilled 1.5 mL Eppendorf tubes and stored for one hour at -20 °C before for RNA isolation. The other two serial sections of each mouse brain were thaw mounted on glass slides as described above. One serial section was immediately used for a 1-hour DESI-MSI image acquisition as detailed below. The last serial section of each mouse brain was left in ambient conditions for 1 hour as a no-DESI-MSI control.

After DESI-MSI, mouse brain sections were removed from the glass slide with a razor blade and placed in 1.5 mL centrifuge tubes. The RNA from all serial sections was extracted using a Qiagen RNeasy Mini Kit following the Qiagen RNeasy protocol (HB-0435). RNA was stored at -80 °C and analyzed using an RNA ScreenTape with RNA ScreenTape sample buffer (Agilent Technologies) on a 2200 TapeStation (Agilent Technologies). RIN scores were calculated using TapeStation Analysis Software v5.1 (Agilent Technologies).

### Analysis of Human Cancer Samples

Prior to DESI-MSI analysis, a section of each tissue was taken for tissue RNA quality analysis. Qiagen RNeasy kits were used to extract RNA from samples and RNA quality was assessed using an Agilent TapeStation with an RNA screen tape to calculate a RIN score as described above. A RIN score greater than 4 was required for subsequent analysis. One section of each tissue was prepared for analysis with the Desium workflow. For 2 samples (BC-1 and LC-2) an adjacent serial section was taken to perform the standard Visium CytAssist alone as a control. A third serial section was taken from BC-1 and analyzed with the Desium workflow because some tissue detachment from the glass slide along the edges of the tissue occurred that resulted in tissue folding while performing the Visium CytAssist portion of the Desium workflow.

### DESI-MSI

After thaw mounting, each sample was allowed to equilibrate for five minutes at room temperature. Slides were imaged on a Xevo G2-XS QTOF (Waters) with a DESI-XS source in negative ion mode. The instrument was placed in sensitivity mode for DESI-MSI acquisition. Solvent spray was 100% methanol with a flow rate of 3 μL/min and nitrogen gas pressure at 1 atm. Samples were analyzed in with a capillary voltage of 0.65 kV. The internal collision gas was set to 6 and the resolution of the instrument was set to 22,000. The geometry of the DESI sprayer was 75 degrees relative to the stage. Images were acquired at 100 x 100 μm^2^ pixel size. The DESI-XS stage rate was set to (200 μm/s), and the instrument scan time was set to 0.458 seconds. Spectral acquisition covered the *m/z* range of 100-1500. Each image took between 30 – 60 minutes to acquire, depending on the size of the tissue.

### H&E Staining and VST CytAssist

Immediately following DESI-MSI image acquisition, tissues were H&E stained with RNase free solvents and imaged with a slide scanner (NanoZoomer, Hamamatsu) following the 10X Genomics H&E staining protocol CG000614 for fresh frozen tissue. Slides were then processed by the Genomic and RNA Profiling Core at Baylor College of Medicine following the 10X Genomics Visium CytAssist protocol CG000495 with no variation. This included probe hybridization, ligation, transfer, extension, library creation, and sequencing. The probe set used was the Visium human transcriptome probe set v2.0, targeting 18,085 genes from the GRch38-2020-A human transcriptome. Visium V4 Slides -FFPE v2 were used for probe capture, and libraries were sequenced with a NovaSeq 6000 sequencer.

### Pathological Evaluation of Tissues

Pathological evaluation of H&E images was performed by C.N. and M.R. board certified breast and lung pathologists, respectively. Overall cancer subtype was confirmed, and histological features were annotated, including tumor cells, normal cells, immune cells, stroma, and necrosis were indicated in each tissue section using the web-based annotation tool included in the Weave platform for Spatial Biology (Aspect Analytics NV, Genk, Belgium).

### MSI and Visium Data Processing

MSI data was corrected for mass drift using the Waters CLMC program with *m/z* 885.5499 as an internal lock mass, threshold counts at 1000, tolerance PPM at 500, correction period at 1 minute, and the combine time at 0.1 minutes. Mass spectra were then centroided in Mass Lynx V4.2 (Waters) using the automatic peak detection tool in negative ion mode. Mass drift corrected, centroided data was converted from Waters Raw files into MZML format using the ProteoWizard MSconvert tool. MZML files were then processed into IMZML file formats using MSIreader V1.03. IMZML imaging data was preprocessed by performing TIC normalization, peak picking, and subsequent binning. Peak picking was performed on the mean mass spectrum across all datasets. Peaks in the mean mass spectrum with an intensity below 0.05% of the base peak were discarded, resulting in a total of 2086 retained peaks. Intensities were binned per spectrum using a 30-ppm window around the selected peaks.

Visium sequencing libraries were processed using SpaceRanger v2.1.1 (10X Genomics). FASTQ files were mapped to the prebuilt GHCh38 human reference genome provided by 10X Genomics. Visium data was further processed in R v4.3.2 using the Seurat v5.0.1 package. Sample gene counts were normalized using the sctransform function from the Seurat package.

### Comparison of sequencing results between Desium and Visium workflows

For samples LC-1 and BC-1, distributions of Genes/Spot, and UMI Counts/Spot were plotted directly with Seurat. The mean UMI Counts/gene for each workflow and sample were exported from Seurat, and a correlation plot comparing these values was generated using ggplot2. Desium and Visium data for LC-1 and BC-1 each were merged into one Seurat object while retaining their original identities. UMAP clustering of each dataset was performed by constructing a shared nearest neighbor graph based on the first 30 principal components (FindNeighbors) and running a uniform manifold approximation and projection (UMAP) embedding also based on the first 30 principal components (RunUMAP); this was followed by graph-based clustering (FindClusters). UMAP space dimension plots and spatial dimension plots were plotted directly from Seurat.

### Spatial multi-omics integration pipeline

MSI and VST data were indirectly registered to each other by their respective H&E-stained microscopy images (i.e., post-MSI H&E image and the Visium CytAssist H&E image). To co-register MSI data to its H&E image, a couple of representative images from MSI were generated by dimensionality reduction methods e.g. uniform manifold approximation and projection (UMAP) and non-negative matrix factorization (NMF), to guide the manual non-rigid co-registration workflow in the Weave platform (Aspect Analytics NV). The post-MSI H&E image was then co-registered to CytAssist Image, whose spatial reference is inherent to VST data, via an automatic linear co-registration workflow from the Weave platform. Finally, MSI data was transformed to the spatial coordinate system of VST data.

Before the granularity matching, each VST data was inspected, and spots that were apart from the main tissue area or overlaying folded tissue regions were removed. Multiple neighboring MSI pixels (100 µm /pixel) were aggregated to a single representative spectrum to match each spot from VST (55 µm in diameter with a 100 µm distance between spots) by a Gaussian weighting algorithm.^17^

### Dimensionality reduction via non-negative matrix factorization

Non-negative matrix factorization (NMF) was applied to MSI and VST data separately due to its non-negativity constraint and ability of interpretable parts-based representation generation. NMF was implemented via the Python package sklearn.decomposition.NMF with Kullback–Leibler divergence (KL-NMF) as the cost function and Multiplicative Update was used as the solver.

### Correlation analysis

Pearson correlation was implemented via Python package numpy.corrcoef. Correlation coefficients range from -1 (negative linear relationship) to 1 (positive linear relationship), with 0 implying no correlation between two datasets. A list of gene to *m/z* value correlations for individual samples above 0.4 and below -0.4 for all *m/z* values including unknown *m/z* values can be found in supplemental table 3. Gene to metabolite correlations for identified metabolites and lipids are found in supplemental table 4. Mean correlations for all m/z values including unknowns above 0.3 or below -0.3 are found in supplemental table 5. Mean correlations for identified metabolites and lipids are found in supplemental table 6.

### Lipid and metabolite annotation/identification

A total of 159 m/z were tentatively identified with high mass accuracy, within 5 ppm of known biological species commonly observed in DESI-MSI experiments. 63 of these *m/*z features were annotated putatively with MS2 including those described in the main text of this manuscript. Both MS1 and MS2 level annotations are found in supplementary table 2. MS2 was performed using direct injection ESI-tandem MS (MS/MS) was performed on lipid and metabolite extracts obtained from the leftover tissue block from Desium analyses. Lipid and metabolite extracts were obtained using the Bligh-Dyer method on ∼10 mg of each tissue (obtained by weighing 50 μm tissue sections of each tissue). Briefly, tissue sections were suspended in 300 μL of ice-cold water and 400 μL of chilled methanol. Suspended tissue sections were homogenized by vortexing vigorously for 30 seconds and sonication in a room temperature water bath for 5 minutes. 800 μL of chloroform was then added to each sample, vortexed again for 30 seconds, and then shaken at room temperature for 20 minutes. Extracts were then centrifuged at 18000 g for 20 minutes at 4 °C. The upper aqueous methanol layer and lower organic chloroform layer were both carefully transferred to separate tubes for each sample and dried at room temperature in a SpeedVac. Samples were reconstituted with 100 μL methanol. 5 μL of each reconstituted sample was injected for direct ESI analysis on an Orbitrap Exploris 240 (Thermo) using 90% methanol/10% water carrier solvent. Targeted MS/MS was performed by using sample specific inclusion lists with a 20-ppm peak isolation window and a 30% normalized HCD collision energy. MS/MS data was processed using Thermo Xcalibur Qual Browser software. MS2 fragment matches for the compounds reported in the main text are found in tables 7-15 of the supplemental information. Lipid names were based on the LipidMaps guidelines on shorthand notation for lipid structures published by Liebisch et al.^18,19^

## Acknowledgments

This study was supported by the Welch Foundation (grant number Q-1895-20220331), the National Cancer Institute of the National Institutes of Health (award number R01CA284742), and The Marcus Foundation. This project was also supported in part by the Genomic and RNA Profiling (GARP) Core at Baylor College of Medicine with funding from the NIH S10 (1S10OD023469), NIH NCI (P30CA125123), and CPRIT (RP200504) grants. We acknowledge the GARP Core for performing Visium CytAssist protocols and subsequent sequencing. We thank Daniel Kraushaar and Patricia Castro for helpful communications about the Visium CytAssist technique. We acknowledge Baruch Berger and Marc Claesen from Aspect Analytics NV for their contributions to the Weave pipeline, which enabled DES data analysis and annotation transfer.

## Author Contributions

L.S.E. and T.M.G. initiated and planned experiments. T.M.G. and Y.S. planned and performed experiments. T.M.G. and F.E.J. processed and analyzed DESI-MSI and VST data and generated figures. W.Z., T.T., N.V., and N.H.P. processed and analyzed DESI-MSI and VST data, performed coregistration, and subsequent statistical analyses. C.N. and M.R. performed pathological evaluation of cancer tissues. T.M.G. and L.S.E. wrote the paper with input and edits from other authors. All authors read and approved the final paper.

---

**Extended Data Figure 1:**
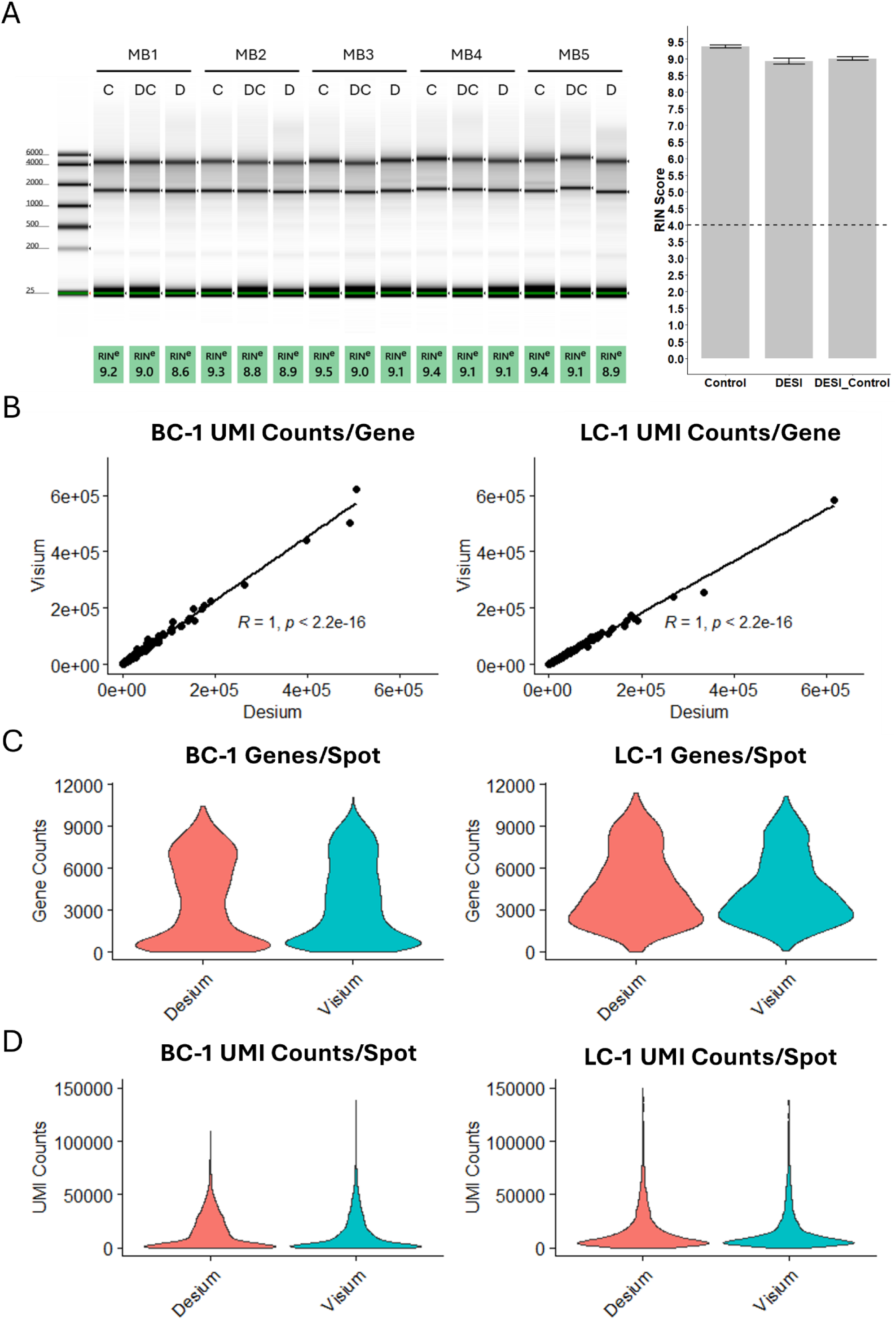
DESI-MSI has almost no effect on tissue RNA Quality. **a.** The effect of DESI-MSI on RNA quality in tissues was evaluated on 5 fresh frozen mouse brains (MB1 – MB5) by comparing unanalyzed controls (C), a no-spray DESI controls left at room temperature for one hour (DC), and tissue analyzed by DESI-MSI with 100% methanol spray for 1 hour (D). RIN scores were calculated for each condition as an indicator of RNA quality. A virtual gel calculated from an Agilent TapeStation and corresponding RIN scores for each condition and are summarized in bar plot. A minimum RIN score of 4 is required to perform the VST CytAssist protocol and is marked with a dotted line. **b.** Unique Molecular Identifier (UMI) count/gene comparison between workflows for breast cancer sample BC-1 and lung cancer sample LC-1 where each spot is represents a single gene and its UMI counts in each condition. R = correlation coefficient. **c.** Distribution of genes/spot between workflows for samples BC-1 and LC-1. **d.** Distribution of UMI counts/spot between workflows for samples BC-1 and LC-1.

**Extended Data Figure 2:**
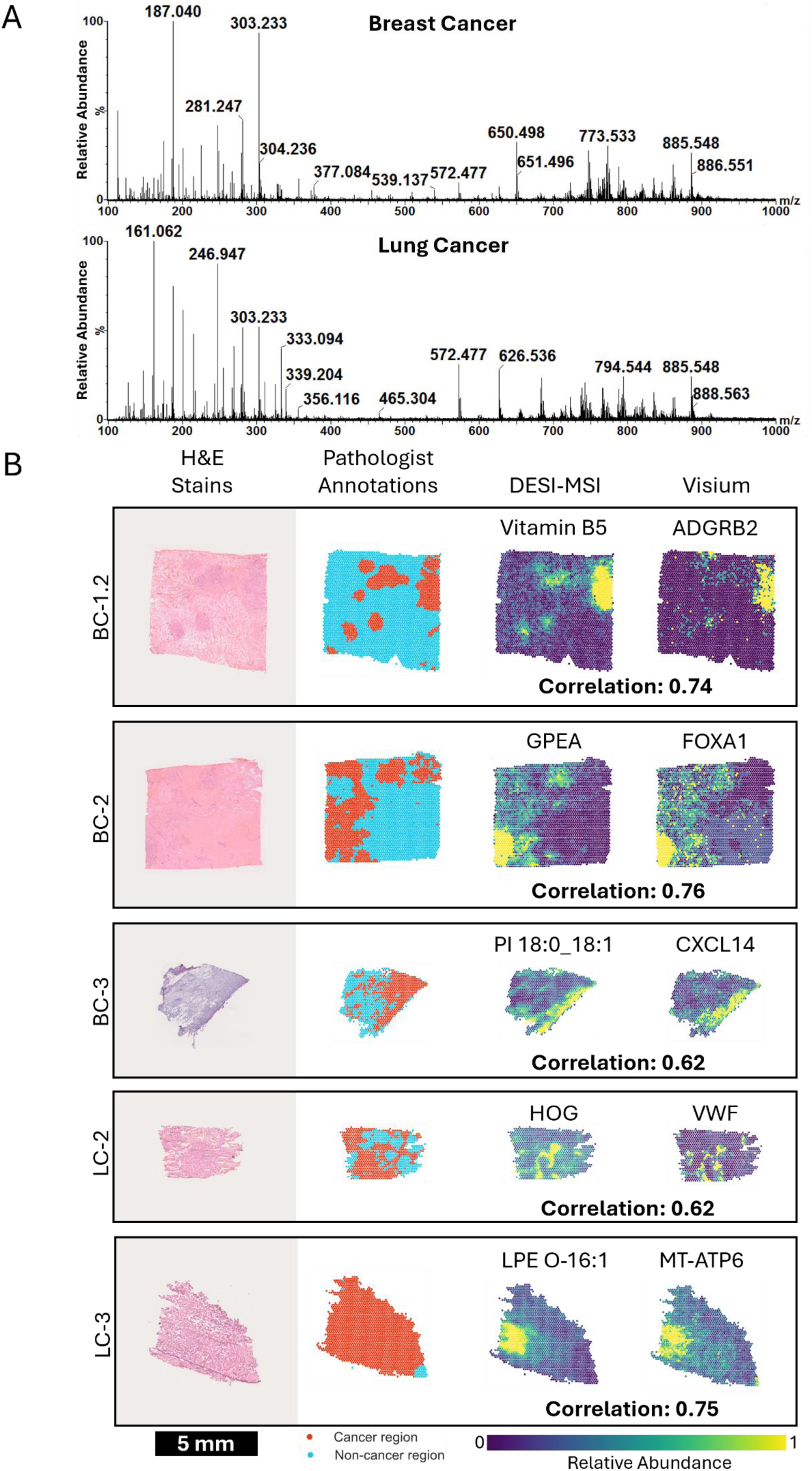
Representative DESI-MSI data and additional gene/metabolite correlations. **a,** Representative breast and lung cancer MS1 spectra acquired with DESI-MSI in negative ion mode during the Desium workflow for samples BC-1 and LC-1. Lipid region *m/z* 500-1000, metabolite/fatty acid region *m/z* 100-500. **b,** A collection of the top individual lipid/metabolite to gene Pearson correlations for breast and lung cancer samples BC-1.2, BC-2, BC-3, LC-2, and LC-3 for which putative MS2 annotations could be obtained. These are aligned with cancer regions identified by a board-certified pathologist for comparison. GPEA: glycerylphosphorylethanolamine, PI: Phosphatidylinositol, HOG: 4-hydroxy-2-oxoglutarate, LPE lysophosphatidylethanolamine. Lipid fatty acid chain notation: (fatty acid chain one)_(fatty acid chain two), (# of carbons in FA chain):(# of double bonds in chain).

**Extended Data Figure 3:**
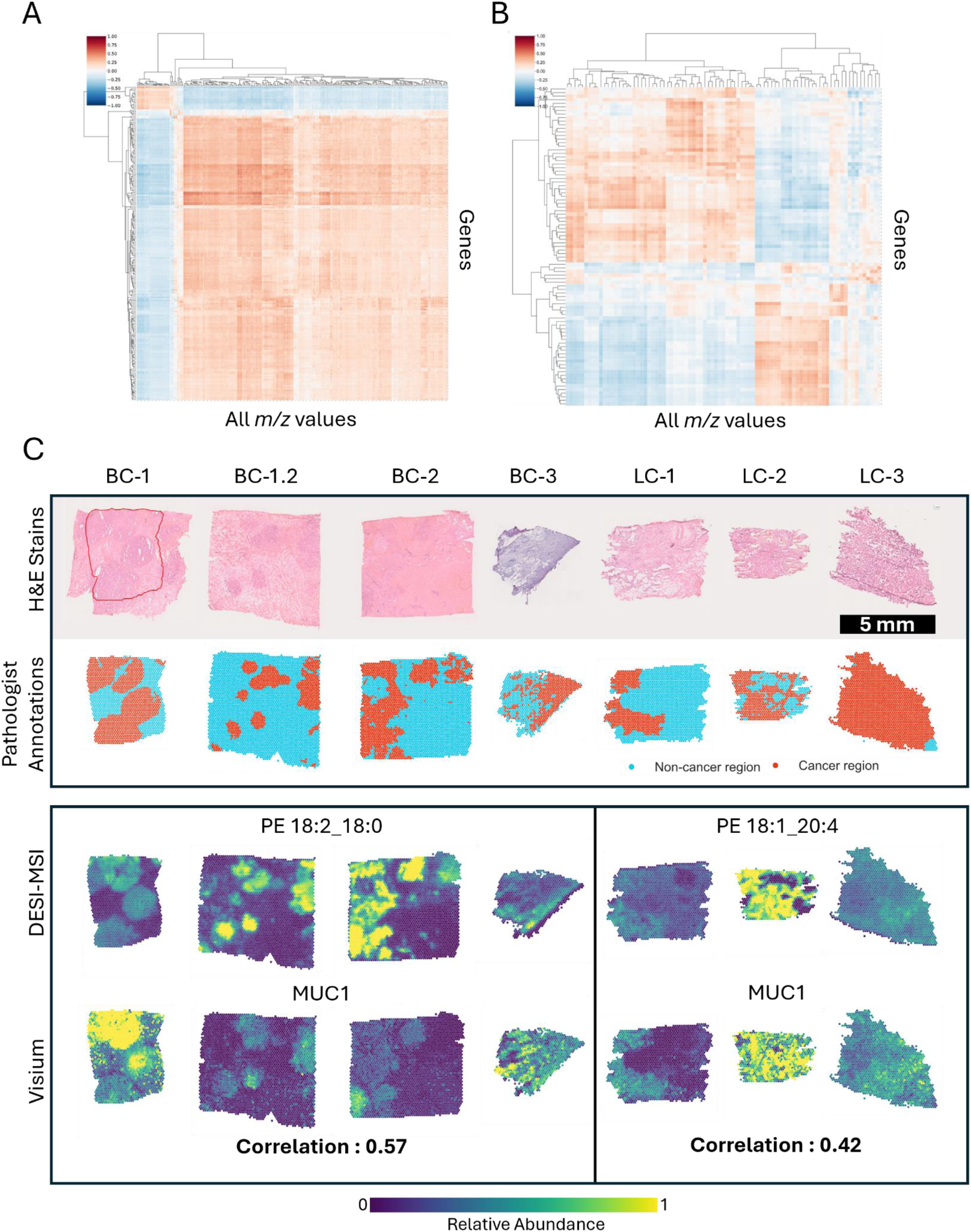
Gene and Lipid Correlations across Tissue Type. **a,** Heatmap showing average Pearson correlations across all breast cancer samples between all *m/z* values (including unidentified peaks) and transcripts. **b,** Heatmap showing average Pearson correlations across all lung cancer samples between all *m/z* values (including unidentified peaks) and transcripts. **c,** The highest top averaged lipid/metabolite to gene Pearson correlations for breast cancer and lung cancer aligned with histological features that pathologists identified as cancer regions. PE: phosphatidylethanolamine. Lipid fatty acid chain notation: (fatty acid chain one)_(fatty acid chain two), (# of carbons in FA chain):(# of double

**Extended Data Figure 4:**
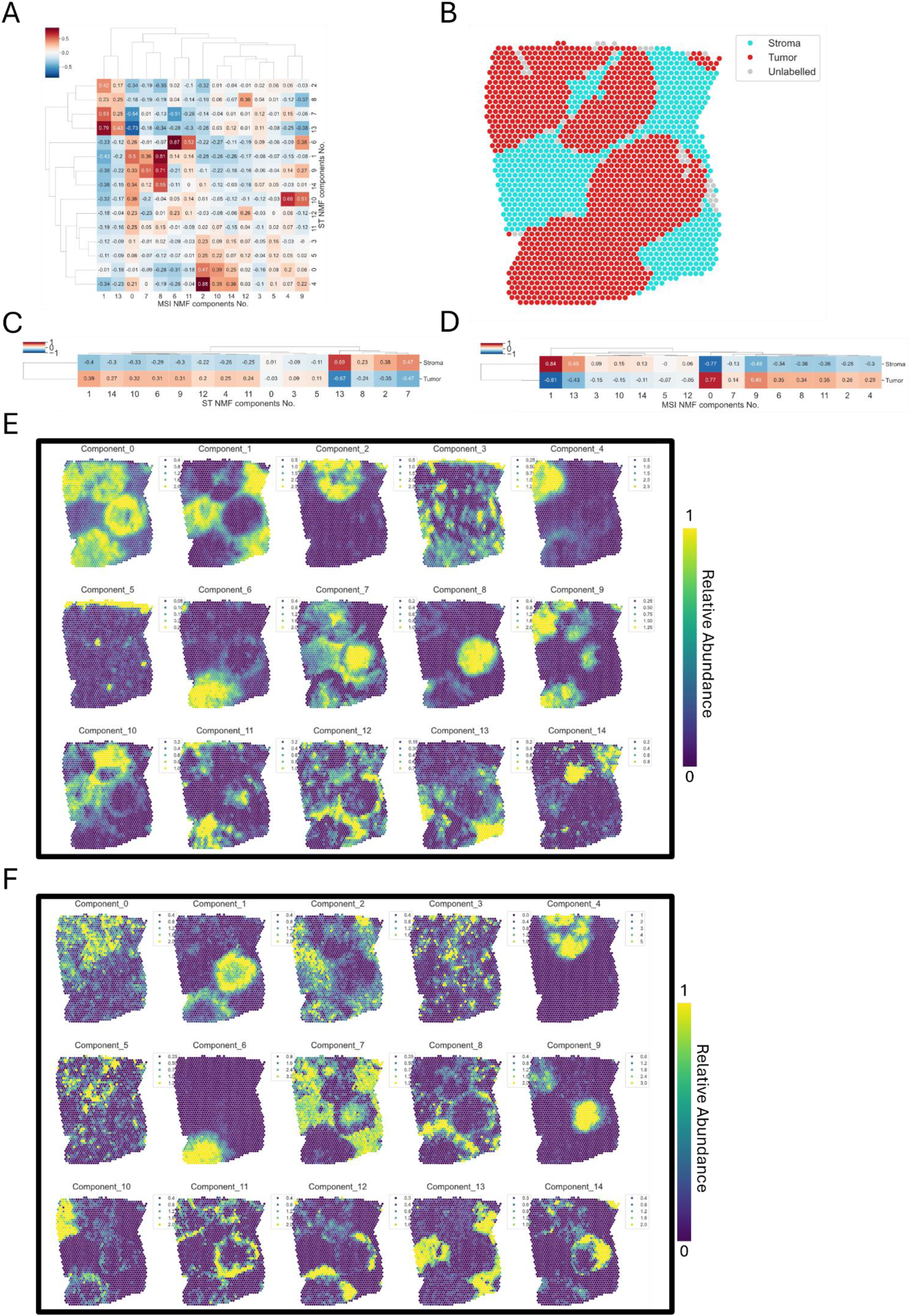
BC-1 Correlations between DESI-MSI and VST NMF components. **a,** Heat map showing Pearson correlation scores between DESI-MSI and VST NMF components for breast cancer sample BC-1 **b,** Pathologist annotations of BC-1 transformed into Visium spot format. **c-d,** Heatmaps showing the Pearson correlation score of each NMF component with pathologist identified regions. **e,** NMF components from the MSI data. **f,** NMF components from the VST data.

**Extended Data Figure 5:**
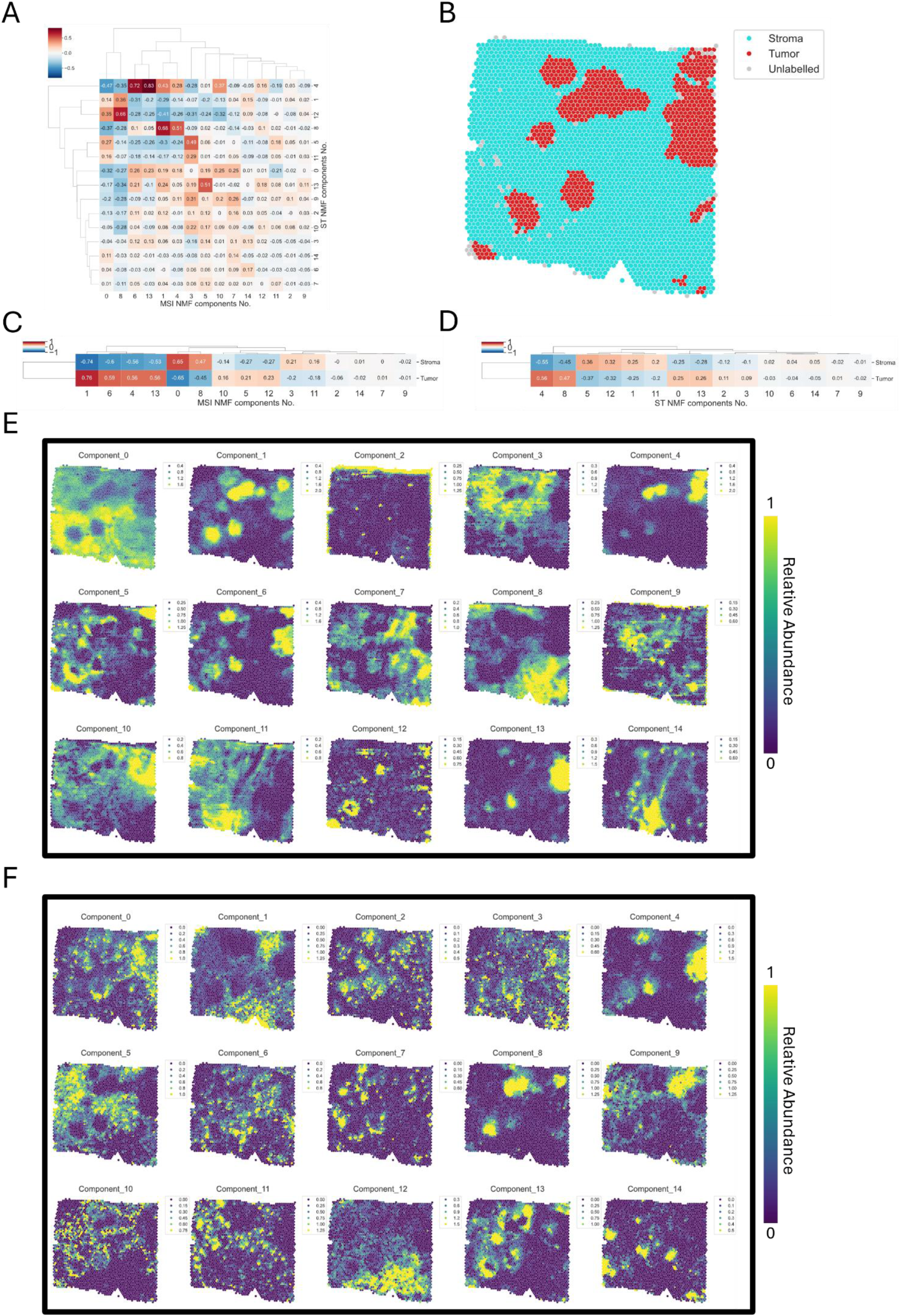
BC-1.2 Correlations between DESI-MSI and VST NMF components. **a,** Heat map showing Pearson correlation scores between DESI-MSI and VST NMF components for breast cancer sample 1.2 **b,** Pathologist annotations of BC-1.2 transformed into Visium spot format. **c-d,** Heatmaps showing the Pearson correlation score of each NMF component with pathologist identified regions. **e,** NMF components from the MSI data. **f,** NMF components from the VST data.

**Extended Data Figure 6:**
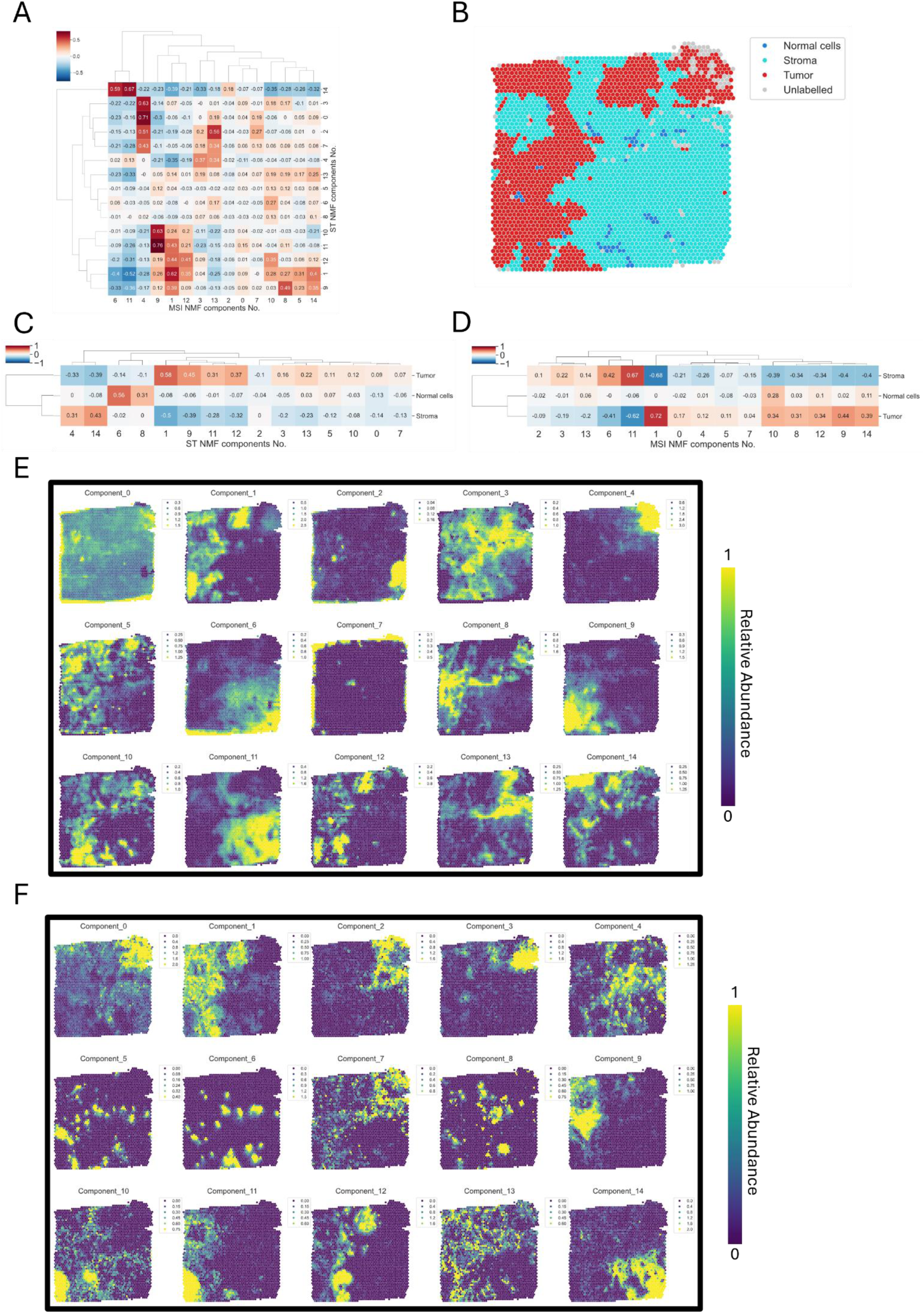
BC-2 Correlations between DESI-MSI and VST NMF components. **a.** Heat map showing Pearson correlation scores between DESI-MSI and VST NMF components for breast cancer sample BC-2 **b.** Pathologist annotations of BC-2 transformed into Visium spot format. **c-d.** Heatmaps showing the Pearson correlation score of each NMF component with pathologist identified regions. **e,** NMF components from the MSI data. **f,** NMF components from the VST data.

**Extended Data Figure 7:**
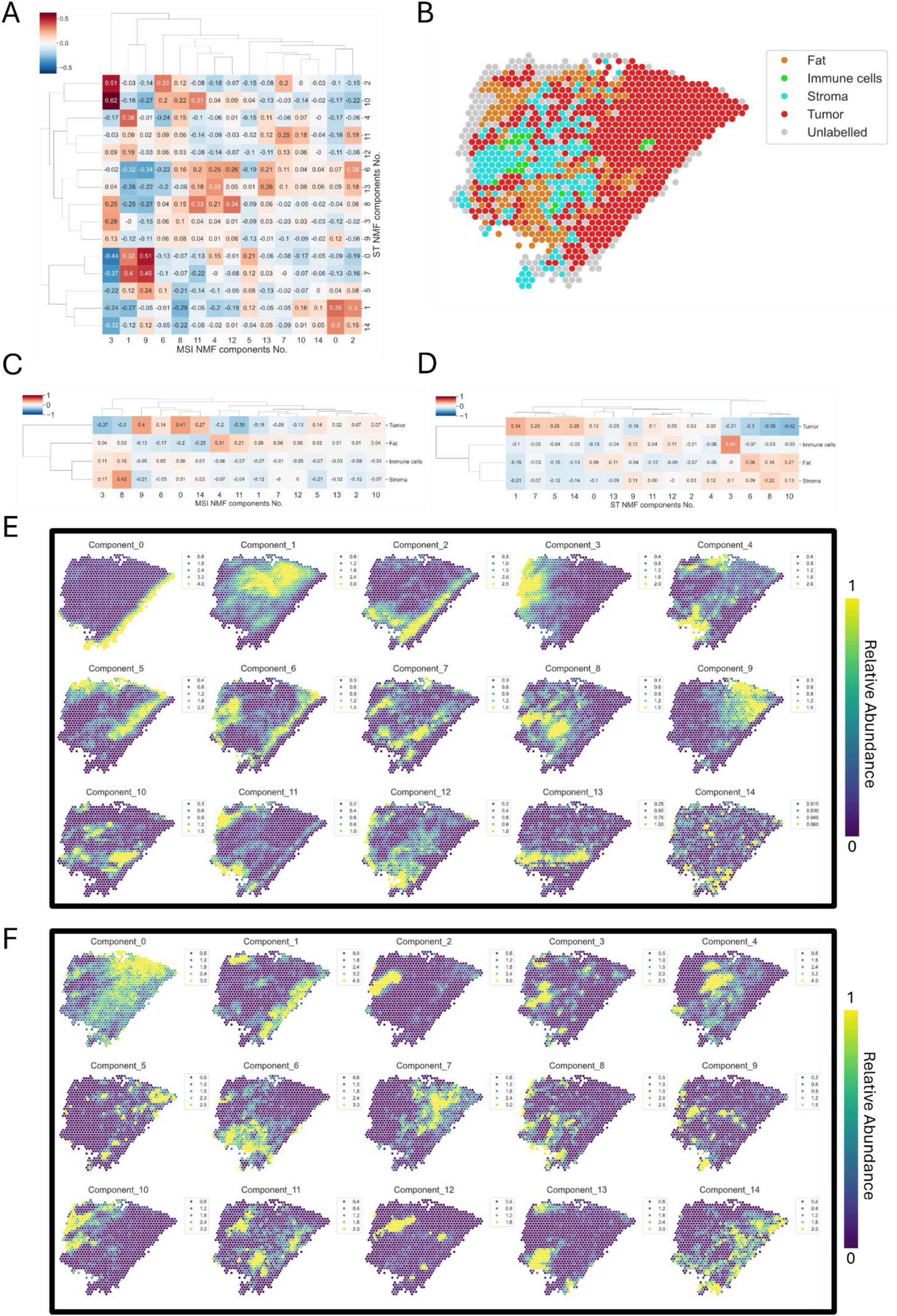
BC-3 Correlations between DESI-MSI and VST NMF components. **a,** Heat map showing Pearson correlation scores between DESI-MSI and VST NMF components for breast cancer sample BC-3 **b**, Pathologist annotations of BC-3 transformed into Visium spot format. **c-d,** Heatmaps showing the Pearson correlation score of each NMF component with pathologist identified regions. **e,** NMF components from the MSI data. **f,** NMF components from the VST data.

**Extended Data Figure 8:**
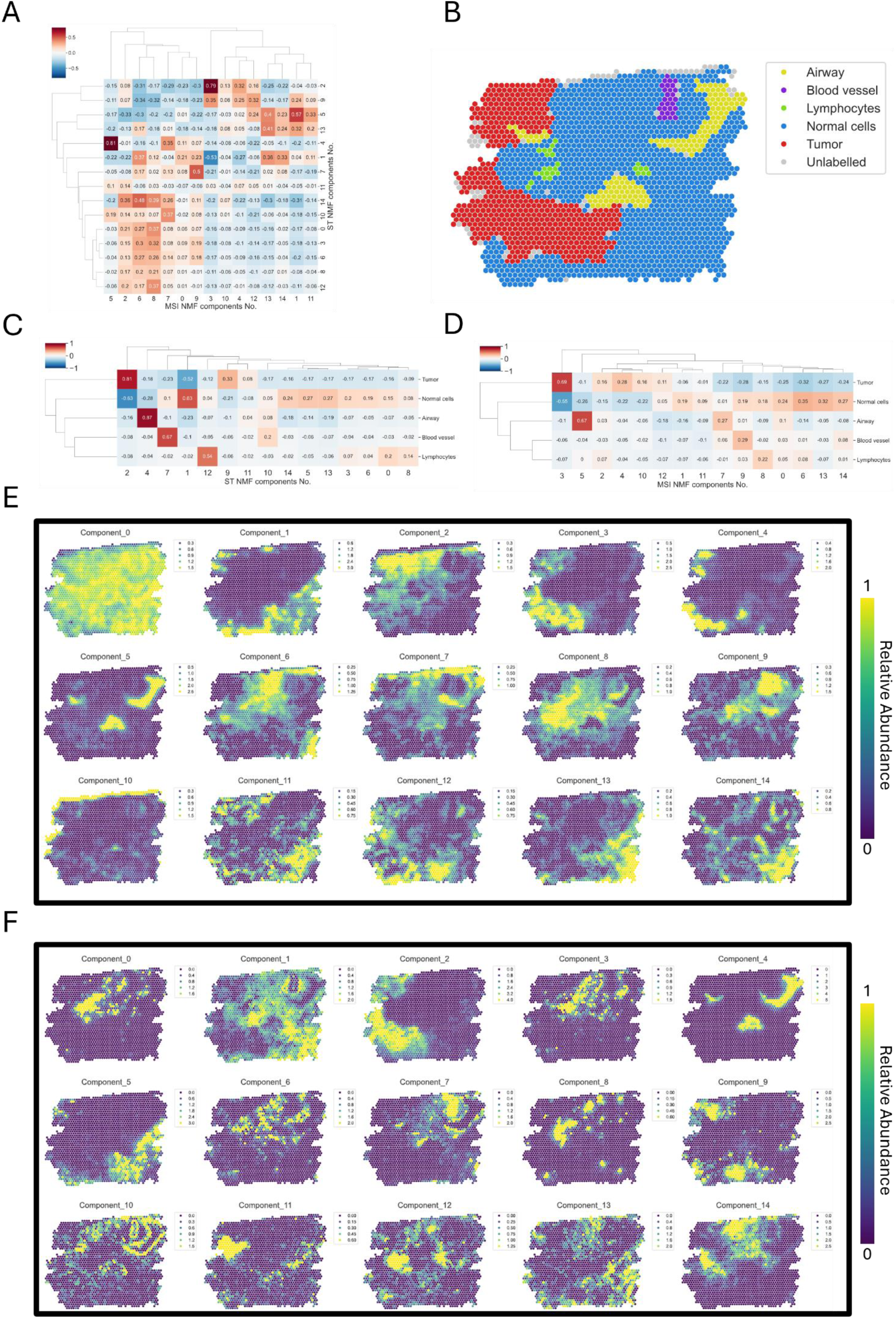
LC-1 Correlations between DESI-MSI and VST NMF components. **a,** Heat map showing Pearson correlation scores between DESI-MSI and VST NMF components for lung cancer sample LC-1 **b**, Pathologist annotations of LC-1 transformed into Visium spot format. **c-d,** Heatmaps showing the Pearson correlation score of each NMF component with pathologist identified regions. **e,** NMF components from the MSI data. **f,** NMF components from the VST data.

**Extended Data Figure 9:**
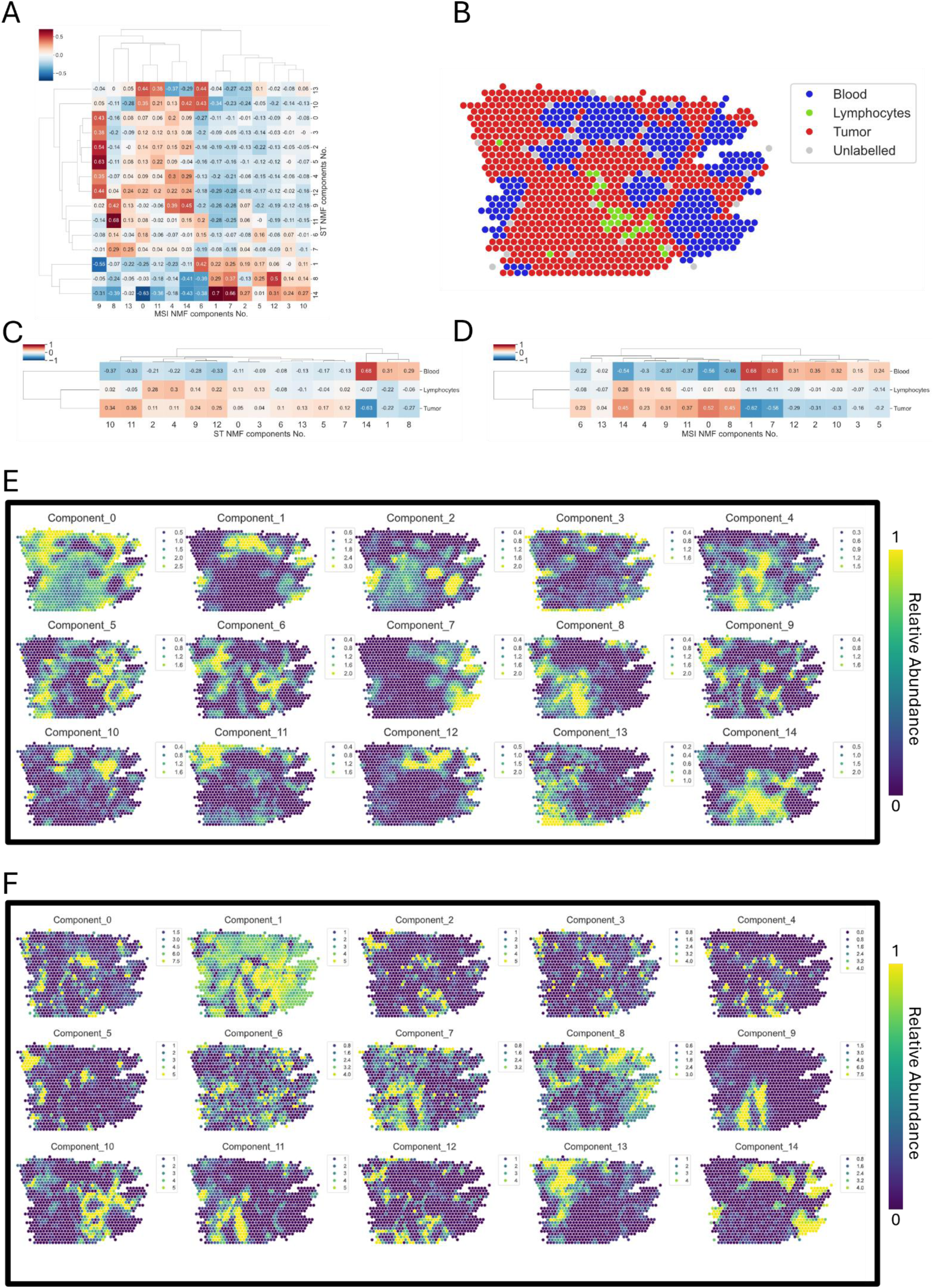
LC-2 Correlations between DESI-MSI and VST NMF components. **a,** Heat map showing Pearson correlation scores between DESI-MSI and VST NMF components for lung cancer sample LC-2 **b,** Pathologist annotations of LC-2 transformed into Visium spot format. **c-d,** Heatmaps showing the Pearson correlation score of each NMF component with pathologist identified regions. **e,** NMF components from the MSI data. **f,** NMF components from the VST data.

**Extended Data Figure 10:**
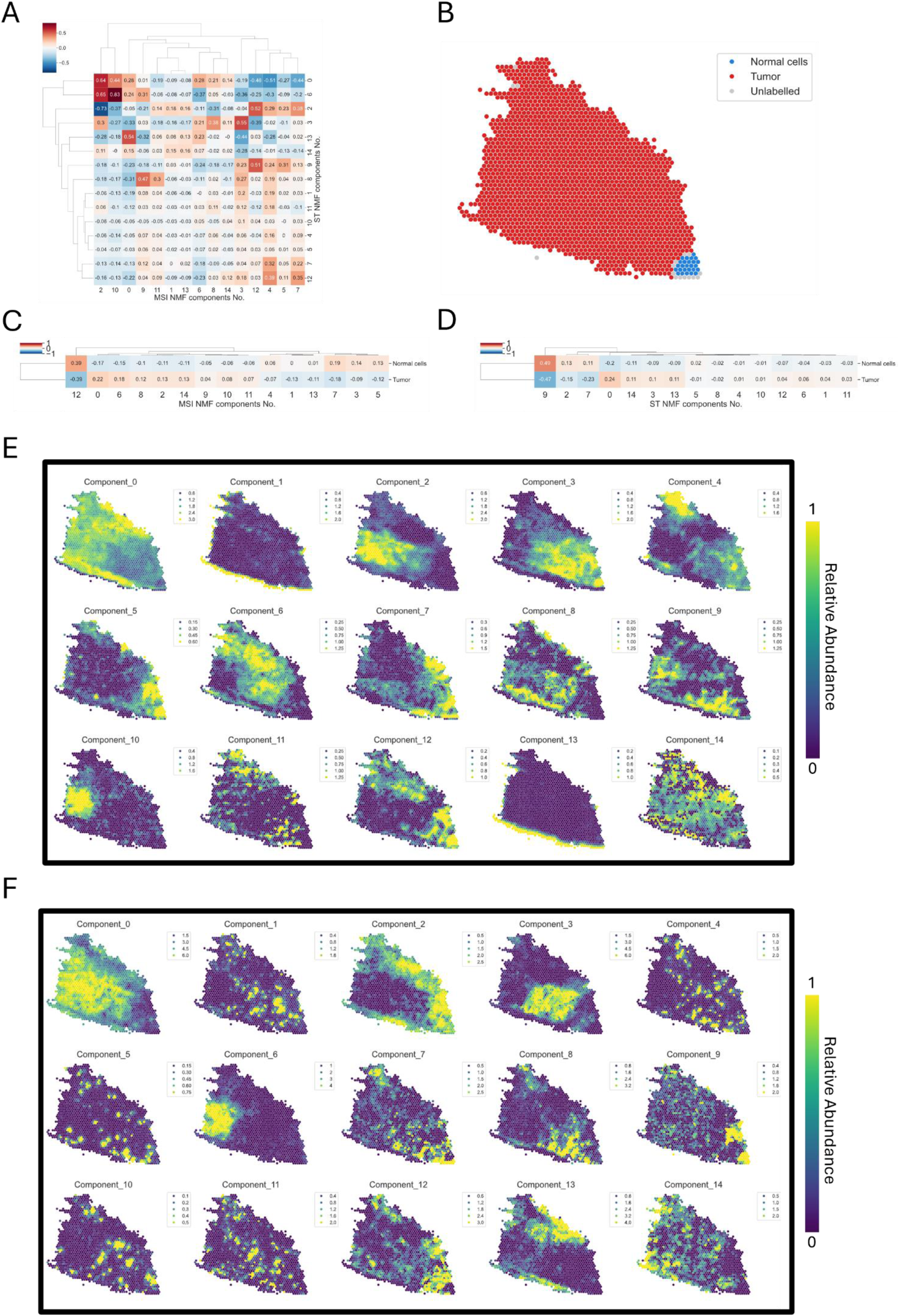
LC-3 Correlations between DESI-MSI and VST NMF components. **a,** Heat map showing Pearson correlation scores between DESI-MSI and VST NMF components for lung cancer sample LC-3 **b,** Pathologist annotations of LC-3 transformed into Visium spot format. **c-d,** Heatmaps showing the Pearson correlation score of each NMF component with pathologist identified regions. **e,** NMF components from the MSI data. **f,** NMF components from the VST data.

